# First study on age and growth of the deep-water Goblin Shark, *Mitsukurina owstoni* (Jordan, 1898)

**DOI:** 10.1101/2020.02.04.934281

**Authors:** Fabio P. Caltabellotta, Zachary A. Siders, Gregor Cailliet, Fabio S. Motta, Otto B. F. Gadig

**Affiliations:** Fisheries and Aquatic Sciences Program, School of Forest Resources and Conservation, University of Florida, Gainesville, FL, USA; Pacific Shark Research Center, Moss Landing Marine Laboratories, Moss Landing, CA, USA; Marine Ecology and Conservation Lab (LABECMar), Federal University of São Paulo, Santos, Brazil; Elasmobranch Lab, Biosciences Institute, Sao Paulo State University, São Vicente, Brazil

**Keywords:** *Mitsukurina owstoni*, Deep-water elasmobranchs, Bayesian age-growth, Goblin Shark

## Abstract

Due to poorly mineralizing structures, ageing deep-water elasmobranchs requires nonconventional techniques. The aim of the present study was to develop a reliable ageing technique using the vertebral centrum to provide information about the age and growth parameters in the Goblin Shark, *Mitsukurina owstoni* (Jordan, 1898). One vertebral centrum from an individual measuring 315.2 cm in total length was analysed. A minimum age of 27 years was estimated. By incorporating priors based on the growth of deep-water sharks and an additional likelihood on *L*_∞_ using data on large male Goblin Sharks, a Bayesian von Bertalanffy growth model was estimated with a male *L*_∞_ of 364 cm total length, weigh 215 kg at *L*_∞_, grow slowly with a *k* equal to 0.049, mature at 16.5 years, and live up to 55 years. Our results are essential to provide useful life history information, with the aim of elucidating the cryptic ecology and biology of this deep-water shark.

## Introduction

Deep-water elasmobranchs comprise nearly half (47.5%) of the known species of sharks and rays of the world (Kyne and Simpfendorfer, 2007). Combined, they represent the highest proportion (57.6%) of species designated as data deficient by the International Union for the Conservation of Nature (IUCN) (IUCN, 2019). The main contributors to the data-deficient status are a lack of life-history information, such as growth rates, age at maturity, and longevity (Kyne and Simpfendorfer, 2010; Dulvy *et al*., 2017).

For the vast majority of deep-water elasmobranchs, poor mineralization of the vertebral centra results in low discernibility in the banding patterns and contributes to the lack of age information (Goldman *et al*., 2012; Cotton *et al*., 2014; Cailliet, 2015). Subsequently, a variety of different structures and alternative ageing methods have been developed: dorsal-fin spines (Irvine *et al*., 2006; Cotton *et al*., 2011), caudal thorns (Henderson *et al*., 2004; Gallagher *et al*., 2006), near-infrared spectroscopy (Rigby *et al*., 2014) and eye lenses (Francis *et al*., 2018). With the deep-water expansion of fisheries, it is imperative to develop life history information for deep-water elasmobranchs and, antecedently, apply ageing techniques to generate critical age-growth information.

The goblin shark *Mitsukurina owstoni* (Jordan, 1898), sole extant member of the family Mitsukurinidae, is one of the largest and most bizarre members of the deep-water elasmobranch world. The species is circumglobal, occurring down to depths of 1000-1500 meters (Kukuev, 1982) and most frequently encountered in coastal submarine canyons. Goblin Sharks reach approximately 410 cm of total length (Compagno, 2001), though there are estimates of larger specimens (Parsons *et al*., 2002). Despite a circumglobal distribution, the species is infrequently observed outside of Sagami Bay, Tokyo Canyon, and Suruga Bay in Japan (Yano *et al*., 2007), and it is considered one of the least known deep-water elasmobranchs with no data available in terms of basic life history information (e.g. age and growth) (Yano *et al*., 2007). In order to better understand the biology, ecology, and conservation status of *M. owstoni*, a reliable ageing technique using the vertebral centrum was developed to provide information about the age and growth parameters.

## Material and methods

### Collection, staining, and ageing

Goblin sharks have been encountered in northern and southern Brazil in the western South Atlantic both through scientific surveys and as a result of a deep-water monkfish, goosefish, and shrimp fishery expansions (Holanda and Asano Filho; Rincon *et al*., 2012, 2017). During an otter trawl operation for deep-water scarlet shrimps on November 27, 2008, one *M. owstoni* specimen was caught (between 700 and 1000 m depth) off the State of Rio de Janeiro, between 23°52’S – 41°52’W and 23°51’S – 42°05’W. The specimen was a mature male weighing 99 kg and measuring 315.2 cm in total length (TL) per Rincon *et al*. (2012). The vertebral column was previously removed from the specimen and kept frozen for about 6 years before attempting to age.

To perform the ageing analysis, vertebrae were manually cleaned and stored in 70% alcohol (Fig. 1A) following standard protocols (Cailliet and Goldman, 2004). Unfortunately, attempts to create bowtie sections were unsuccessful due to the poor mineralization of the intermedialia. Therefore, vertebral centra was first sagittally sectioned (Fig. 1B) along the focus resulting in two bowtie-shaped halves (Fig. 1C) and then the halves were transversally sectioned (Fig. 1B), resulting in four quadrants (Fig. 1D). Each quadrant was stained using the alcian blue staining method adapted from Song and Parenti (1995) (Fig. 1E). This method consisted of using 10 mg alcian blue powder, 20 ml absolute acetic acid and 80 ml 95% ethyl alcohol. Each vertebral quadrant was soaked for 10 minutes intervals then removed, washed with 70% ethyl alcohol, allowed to dry for 30 minutes, then re-submerged and repeated four times (a total of 40 minutes of soaking). At the end of the staining process, one vertebra quadrant was photographed using Motic Image Plus 2.0 imaging software (Motic China Group Co., Ltd.; http://www.motic.com) under transmitted light using a stereomicroscope at a magnification of 10–20x. Three independent readers analyzed the resulting images. Band-pairs, along the corpus calcareum, consisting of one wide band (opaque) and one narrow band (translucent) were identified following the description and terminology detailed in Cailliet et al. (2006) (Fig. 1F). The birthmark was identified as the first distinct band after the focus associated with a slight change in the angle of the corpus calcareum (Fig. 1F).

**Fig. 1:**
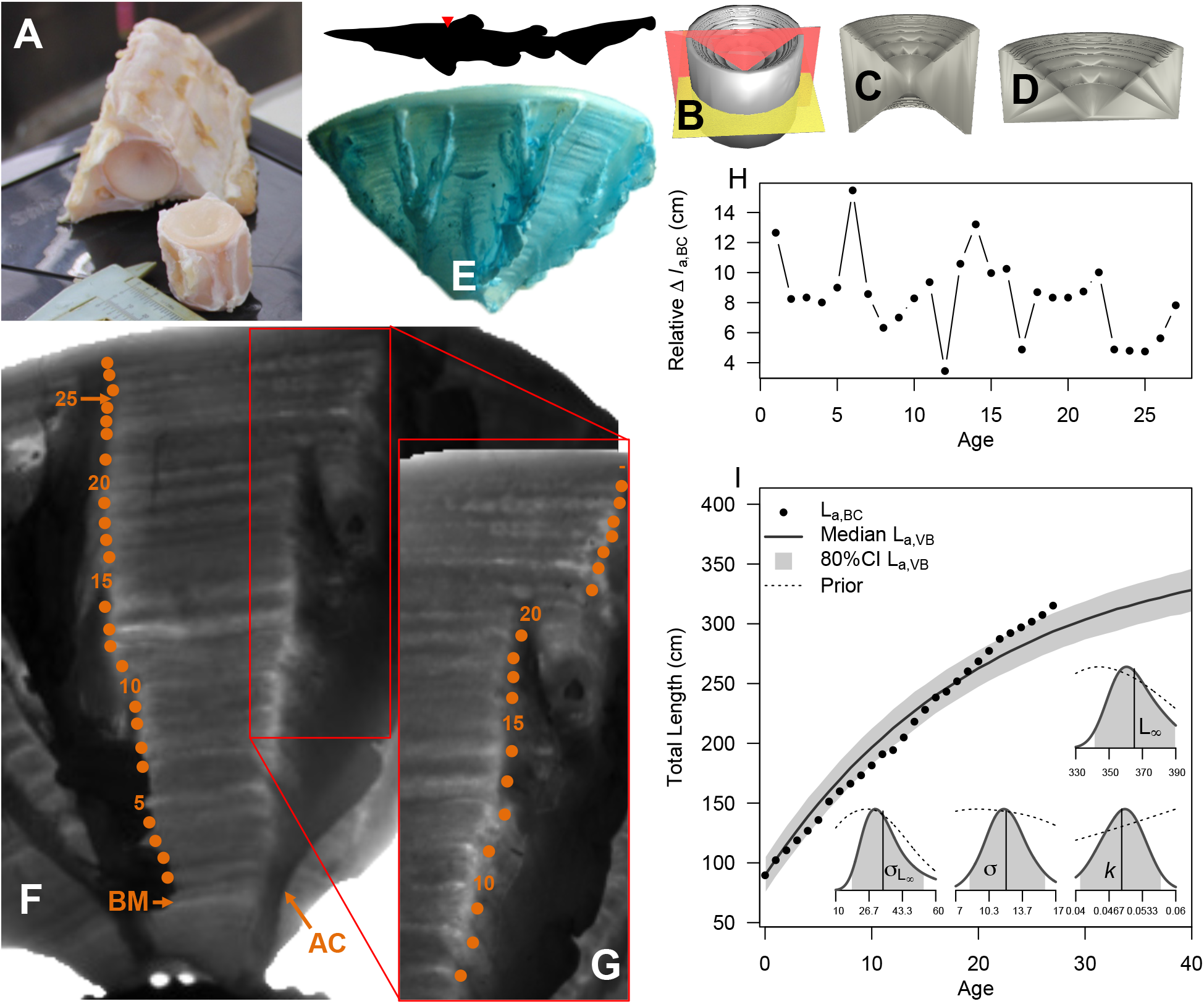
Schematic of the ageing of the vertebrae of *Mitsukurina owstoni* from extraction (A) where the triangle in the silhouette indicates extraction location, sagittal and transversal sectioning (B), the resulting vertebral halves (C) and quadrants (D), to staining with alcian blue mixture (E). The resulting birth mark (BM), angle change (AC), and the median reader count of the number of bands (28 bands) were denoted on the corpus calcareum (F) and in a zoomed in section of the corpus calcareum (G). Using the relative band distance from the BM, back-calculation lengths at age were estimated and the growth increment calculated (H). A von Bertalanffy growth model (I) was estimated based on the back-calculated (points). The median (solid line) and 80% credible intervals (gray polygon) are shown for the growth model predictions as well as for the model’s posterior parameter estimates along with the prior (dashed line).

### Relative Growth Increment

The relative distance from the birthmark, *D_DM_*, and each translucent band, *D_t_*, from 0 (BM) … *A* ages was measured using Adobe Photoshop CC (Adobe Inc., San Jose, California, USA) and used to calculate the relative growth increment between each band (*R_t_*, Equation 1).

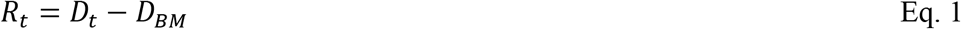

The relative growth increment was used to back-calculate the length at age, *l_t,BC_*, using the Fraser-Lee method (Francis, 1990) (Eq. 2).

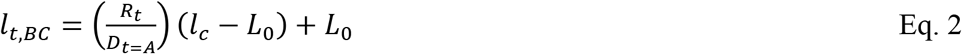

where *l_c_* is the length of capture and *L*_0_ is the length at birth (note that *L*_0_ is substituted for the intercept of the vertebral radius and *l_c_* as this was inestimable with a single specimen). We assumed that length at birth was the mean between the two smallest male and female known specimens, 81.7 cm and 97.5 cm TL respectively (Yano *et al*., 2007; Castro, 2011).

### von Bertalanffy growth modeling

With only the back-calculated length at age data from one specimen, it was necessary to structure the von Bertalanffy growth model (von Bertalanffy 1934)(VBGM) with strong priors, additional data, and following the *L*_0_ formulation (Cailliet *et al*., 2006) (Eq. 3).

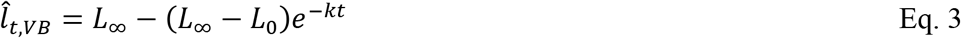

where 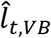 is the predicted length at age from the VBGM, *L*_∞_ is the asymptotic maximum length, and *k* is the Brody growth coefficient. The likelihood of the VBGM component was specified as normal with a mean equal to the predicted length at age (Eq. 3) and a standard deviation of *σ* (Eq. 4).

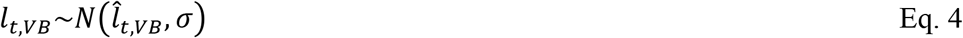

To better inform *L*_∞_, a literature search of the total lengths of large males, greater than *l_c_* (length of capture), was conducted: 315.2 cm (Rincon *et al*., 2012), 317 cm (Yano *et al*., 2007), 320 cm (Cadenat and Blanche, 1981), 320 cm (Ugoretz and Siegel, 1999), 322 cm (Piotrovskiy and Prut’ko, 1980), 355 cm (Masai, 1973), 366 cm (Yanagisawa, 1991), 367 cm (Kurata, 1967), 384 cm (Stevens and Paxton, 1985), and 390 cm (Noden, 1984) [there is 349 cm one specimen that was not included as recent literature has noted the specimen was female not male as initially reported]. The *l_c_* of this study’s specimen is 79% of the length of the largest confirmed male specimen. Using the data available on large confirmed males, *l*_max_, a separate likelihood was used to constrain estimates of *L*_∞_ (Eq. 5).

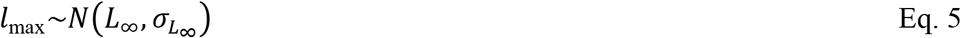

Informative priors were put on the parameters of the VBGM. A normal distribution with a mean of large confirmed male specimens and a standard deviation of 35 was used to specify the prior for *L*_∞_. Based on a study of deep-water elasmobranchs by Rigby and Simpfendorfer (2015), a truncated normal distribution with a mean of 0.1 and a standard deviation of 0.044 was used to specify a prior for *k*. The VBGM was implemented in a Bayesian framework in JAGS (Plummer, 2003) using the *jagsUI* package (Kellner, 2018) in *R* (R Core Team, 2018). Due to JAGS using normal priors based on precision rather than a standard deviation, a truncated normal distribution with mean of 10 and a standard deviation of 10 was used to specify the prior for *τ*, where 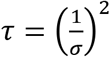. A truncated normal distribution with a mean of 25 and a standard deviation of 20 was used to specify the prior for precision parameter *τ*_*L*∞_. From the VBGM parameters, the theoretical age at length zero (*t*_0_), age at maturity 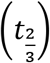 (Beverton and Holt, 1959), and longevity (*t*_0.95_) (Taylor, 1958; Ricker, 1979) were derived by rearranging the VBGM in terms of half-lives (Cailliet *et al*., 2006)(Eq. 6). Mortality at age one, *M*_*t* = 1_, was derived from the VBGM parameters as shown in Charnov *et al*. (2013) (Eq. 7).

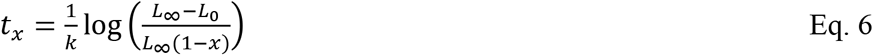

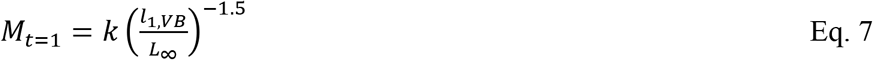

Models were run over four chains with 2,500 samples for adaptation, 5,000 for burn-in, and 1,250 kept samples per chain at a thinning rate of 100 (132,500 total samples per chain). Models were checked for convergence visually and confirmed if the potential scale reduction factor, 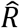, was less than 1.1 (Gelman and Rubin, 1992). During the sampling a posterior predictive check was done to derive model posteriors of the von Bertalanffy length at age, *l_t,VB_*. All data and code is provided at Open Science Framework (DOI 10.17605/OSF.IO/C2ZSW; https://osf.io/c2zsw/?view_only=e5df616cd53a482db0b6d56e77f0ef4c).

## Results

Age readings resulting from the three independent readers were: 27, 28 and 30 years (Fig.1F-G). The growth increment from the back-calculated length at age, *l_t,BC_*, ranged from 3.4 to 15.4 cm with an mean annual growth rate of 8.4 cm. Growth was spasmodic with pulses of rapid growth in ages 5-6 and 14 with pulses of reduced growth in ages 12, 17, and 23-27 (Fig. 1H). The von Bertalanffy growth model converged with all 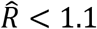. The model predicted greater median lengths at age, *l_t,VB_*, for ages 3-18 and lower lengths at age for ages 19-27 than predicted from the back-calculation, *l_t,BC_* (Fig. 1I). The median parameter estimates of the von Bertalanffy growth curve were 360 cm (*L*_∞_), 0.0508 years^−1^ (*k*), 12.5 (*σ*), and 30 (*σ*_*L*∞_) (Table 1). Median parameter estimates of the derived life history characteristics were −5.64 years (*t*_0_), 16 years 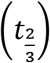, 53.4 years (*t*_0.95_), and 0.0159 (*M*_*t*=1_) (Table 1).

**Table 1:** Posterior estimates of the median and 80% credible intervals (in parentheses) for the von Bertalanffy growth parameters and derived life history parameters.

| Parameter | Median |
| --- | --- |
| $L_{\infty}$ (cm) | 364 (350 – 392) |
| $k$ (years<sup>-1</sup>) | 0.0494 (0.0427 – 0.0544) |
| $\sigma$ | 12.1 (9.61 – 14.9) |
| $\sigma_{L_{\infty}}$ | 33.33 (23.8 – 55.2) |
| $t_0$ (years) | -5.73 (-6.11 – -5.4) |
| $t_{\frac{2}{3}}$ (years, maturity) | 16.5 (14.8 – 19.7) |
| $t_{0.95}$ (years, longevity) | 54.9 (49.7 – 64.2) |
| $M_{a=1}$ | 0.0154 (0.0125 – 0.0193) |

Based on the derived life history characteristics, Goblin Sharks are long-lived, late maturing, and have low natural mortality. Intriguingly, the annual growth increment stays at or below the mean growth increment for our specimen following the estimated age at maturity. A logarithmic regression based on Rigby and Simpfendorfer (2015) for deep-water elasmobranchs (*k*_males_ = −0.067 log(*S_max_*) + 0.586; *R*^2^ = 0.096; *F* = 5.752, *p* = 0.02), estimated a lower *k* of 0.036 based on the median *L*_∞_ of our VBGM than the *k* the VBGM estimated.

## Discussion

In the present study, we describe the first age and growth parameters of *M. owstoni* using the staining technique with alcian blue to reveal the classic opaque-translucent banding, seen in more strongly mineralizing structures. Despite the rapid advancements in ageing poorly mineralizing structures of some deep-water sharks species, no prior information is available about the age and growth, let alone the biology, of Goblin Sharks (Finucci and Duffy, 2018). Here, it is shown that males typically grow to about 360 cm in length, weigh approximately 204 kg per the length-weight relationship in Yano et al. (2007), and grow incredibly slow relative to other sharks.

Based on Rigby and Simpfendorfer (2015), only 16% of aged shark populations > 350 cm in maximum size have a *k* lower than the estimated *k* for *M. owstoni*. None of these aged, large-sharks were deep-water species and the most similar species among the large-shark subset, in maximum size and *k*, was Eastern Central Pacific Shortfin Mako, *Isurus oxyrinchus*. Among deep-water elasmobranchs, the most similar species to *M. owstoni* is the Bird-beaked Dogfish, *Deania calcea*, whose maximum size is approximately one-third of the *L*_∞_ estimated for Goblin Sharks. Interestingly, *D. calcea* has a very similar body shape to *M. owstoni* with an elongated rostrum, deep belly, large epicaudal lobe, and a caudal fin with a low aspect ratio.

The Bayesian technique described above allowed the inducement of the first growth curve for *M. owstoni*. Despite that only one specimen has been aged, growth estimates could likely change if more specimens are aged in the future. Nonetheless, by using strong priors on *k*, fixing *L*_∞_, and incorporating additional data on maximum size a relatively robust von Bertalanffy growth model was estimable and relatively informed derived life history characteristics were calculated. Prior to inclusion of the likelihood constraining *L∞*, posterior estimates of *L*_∞_ were often well above the maximum observed male specimen and occasionally greater than the maximum estimate for the species (Parsons *et al*., 2002). In this scenario and given the tight correlation with *L*_∞_ and *k*, the resulting posterior estimates of *k* were approximately zero. This model improvement phenomenon is expected, as Cailliet *et al*. (2006) astutely predicted as well as from a Bayesian viewpoint, and has the potential to be a relatively simple way of improving age-growth estimations (Caltabellotta *et al*., 2019).

Only 14 of the currently known deep-water shark species have been aged (Kyne and Simpfendorfer, 2010). The alcian blue staining technique is another in the growing toolkit for ageing poorly mineralizing structures of deep-water elasmobranchs. For *M. owstoni*, it proved to result in good resolution for determining banding patterns and estimating an age of the specimen. As Cailliet and Goldman (2004) noted the sagittal-transverse sectioning of the vertebrae caused some distortion near the edge of the corpus calcareum resulting in some disparity in the ageing by independent readers (Fig. 1G). The spasmodic growth increment dynamics we observed are intriguing as they might relate to environmentally driven food availability or spatial ontogeny or maturity.

In conclusion, this study estimates that Goblin Sharks exhibit slow growth, late maturity, long life, and low mortality that, in aggregate, is likely to result in greater vulnerability to overfishing (Stevens *et al*., 2000). Fortunately, there are few fishery interactions as Finucci and Duffy (2018) assessed for the recurrent IUCN Red List status of Least Concern. However, as Yano *et al*. (2007) demonstrated significant numbers can be caught if targeted. With the difficulty of ageing deep-water elasmobranchs, alcian blue staining may be another technique to try on species where traditional techniques fail. In addition, the Bayesian VBGM is a way to estimate growth models with few aged specimens by incorporating strong priors and additional data to constrain the estimation of parameters. These data-deficient quantitative tools and new ageing techniques are essential to provide useful life history information and elucidate the cryptic ecology and biology of deep-water elasmobranchs.

## Acknowledgements

This research did not receive any specific funding

